# Uniform resolution of compact identifiers for biomedical data

**DOI:** 10.1101/101279

**Authors:** Sarala M. Wimalaratne, Nick Juty, John Kunze, Greg Janée, Julie A. McMurry, Niall Beard, Rafael Jimenez, Jeffrey S. Grethe, Henning Hermjakob, Maryann E. Martone, Tim Clark

**Affiliations:** European Molecular Biology Laboratory, European Bioinformatics Institute (EMBL-EBI), Wellcome Trust Genome Campus, Hinxton, Cambridgeshire UK.; University of Manchester, Oxford Road, Manchester UK.; California Digital Library, University of California, Oakland CA, USA.; Oregon Health and Science University, Portland OR, USA; ELIXIR, Wellcome Trust Genome Campus, Hinxton, Cambridgeshire UK; University of California San Diego, La Jolla CA, USA; Massachusetts General Hospital, Boston MA, USA; Harvard Medical School, Boston MA, USA

## Abstract

Most biomedical data repositories issue locally-unique accessions numbers, but do not provide globally unique, machine-resolvable, persistent identifiers for their datasets, as required by publishers wishing to implement data citation in accordance with widely accepted principles. Local accessions may however be prefixed with a namespace identifier, providing global uniqueness. Such “compact identifiers” have been widely used in biomedical informatics to support global resource identification with local identifier assignment.

We report here on our project to provide robust support for machine-resolvable, persistent compact identifiers in biomedical data citation, by harmonizing the Identifiers.org and N2T.net (Name-To-Thing) meta-resolvers and extending their capabilities. Identifiers.org services hosted at the European Molecular Biology Laboratory – European Bioinformatics Institute (EMBL-EBI), and N2T.net services hosted at the California Digital Library (CDL), can now resolve any given identifier from over 600 source databases to its original source on the Web, using a common registry of prefix-based redirection rules.

We believe these services will be of significant help to publishers and others implementing persistent, machine-resolvable citation of research data.

## Introduction

Science policy bodies such as CODATA, the Royal Society and the U.S. National Academies have shown significant concern over the past decade about the reliability of published scientific findings and the reusability of research data^1–3^. Policy concerns have been echoed by major funders and by recent statistical and bibliometric research^4–9^. This situation led in 2013-2014 to the development and publication of the Joint Declaration of Data Citation Principles (JDDCP)^10^, which has been subsequently endorsed by over 100 scholarly organizations. The JDDCP outlines core principles on purpose, function and attributes of data citations, the first of which is that data should be considered legitimate, citable products of research^11^.

The JDDCP require that data become a first-class research object that is archived in persistent stores and is cited just as publications are cited, in all cases where: (1) findings or claims are based on the authors’ primary research data; or (2) data from other sources is input to the authors’ analysis.

Implementation of the JDDCP poses a set of interlocking requirements affecting how archived and cited datasets are to be identified; how identifiers are to be managed; to what and how the identifiers resolve; what dataset metadata are required and how they are serialized; and how data citations are to be represented in the final publication. These requirements affect archival data repositories, scholarly publishers, and various infrastructure providers including identifier registries and bibliographic reference managers; and in the long run, such bodies as promotion committees, who will provide the incentives for authors to change their practices.

Our goal in this article is to present a focused solution to the identifiers problem, targeted specifically to technical implementation of *biomedical data citation*. The project on which we report, was one element of a larger effort – the Data Citation Implementation Pilot (DCIP) - which convened Expert Groups for Identifiers, Repositories, and Publishers. These groups communicated with each other to coordinate their efforts but have reported independently on their results.

JDDCP Principle 3 requires that wherever research findings are based upon data, that data be cited. Principle 4 requires that cited, archived data receive a *globally unique, machine-resolvable persistent identifier* that appears in the citing article’s reference list. This is intended not only to help humans locate data, but to facilitate the development of next-generation mashup tools in an ecosystem based on software agents and searchable research data indexes such as DataMed^12^ and OmicsDI^13^.

- *Globally unique and machine resolvable* identifiers must resolve on the Web as HTTP Uniform Resource Identifiers (URIs).
- *Persistent* identifiers must be robust over time to changes in the underlying location of the data.

A reasonable solution to the identifiers problem is to assign Digital Object Identifiers (DOIs) to identify datasets. DOIs are already widely used in the publishing world as persistent identifiers for scholarly publications. They have been adopted by generalist data repositories such as Dryad, FigShare, Zenodo and Dataverse, as well as by domain data repositories outside of biomedicine. Handles^14^, which underlie the DOI system, may also be used directly. The DataCite consortium provides a robust central means for assigning DOIs to data.

However, DOIs are not commonly used for biomedical data, which is partitioned across over 600 autonomous repositories that are independently funded. Instead, in biomedicine there has been a longstanding practice of employing a prefix plus locally-assigned accession number as a unique identifier. It is not clear that creating DOIs for the billions of existing entities now identified via the prefix:accession model would be worthwhile. Even if it did become socially acceptable, financially tractable, and technically achievable, any tangible benefits could be seriously diminished by the complexity and cost of mapping original identifiers to the new DOIs.

To reduce social and technical barriers to data citation at scale, we developed an approach which is adapted from present practice but nevertheless robust, reliable long-term, and compatible with the JDDCP. The life sciences community typically references data entities using their locally-assigned database accession numbers, often contextualizing these local unique identifiers (LUIs) with a prefix corresponding to the assigning authority to clearly indicate the identifier’s repository context.

These may be rendered web-resolvable through subsequent incorporation into a durable HTTP URI. The assignment of a *formal*, *registered* namespace prefix avoids identifier collisions, and incorporation into an access URI presents the prefix-identifier combination to a resolver system using web protocols.

In some subdomains such as ontologies, internally-assigned prefixes may be an integral part of the identifier minting process and IDs always appear in their prefixed form^15^, for instance, GO:0006915.

Concatenating a repository-identifying namespace prefix, a colon, and an LUI (<prefix>:<LUI>), forms what we call a “compact identifier”. This longstanding and broadly adopted approach is used in International Standard Book Numbers (isbn:<LUI>), Digital Object Identifiers (doi:<LUI>) and other classes of identifier with autonomous assignment.

A core component of persistent identification is redirection, without which it is challenging to provide stable identifiers robust against changes in underlying resource provision. It is thus common practice for bioinformatics data repositories to maintain their own locally-scoped resolvers. A “meta-resolver” is a web server that can recognize enough about an incoming URL to properly redirect to collection-specific resolvers, based on the assigning authority specified by the incoming URL. Meta-resolvers therefore provide a single host from which to launch URLs containing compact identifiers, by appending the prefix plus LUI to the URL base name of the meta-resolver.

Meta-resolvers provide resolution of a common identifier syntax, shield people from the details of and changes to individual resolver access methods, and make it easier to associate compact citations with actionable HTTP links; examples are shown below.

Identifiers.org is a type specimen of this class of service, providing several additional services beyond redirection^16^. It provides stable, persistent and resolvable URIs for the identification of life science (and other) data. These are derived using information stored in an underlying Registry, which contains high quality, manually curated information on over 600 data collections, including a unique namespace prefix, a description of the data collection, identifier pattern, alternative URI schemes and a list of hosting resources or resolving locations. When a compact identifier is presented to the resolver, it is redirected to a resource provider, taking into consideration information such as the uptime and reliability of all available hosting resources. The new implementation described here also enables users to resolve to specific resources by using provider codes.

N2T.net is another such service, with redirection targets based on both individual identifier lookup and prefix-based lookup. N2T was first launched in 2006 as a global resolver for Archival Resource Key (ARK) identifiers. Although assigning ARKs requires neither fees nor a particular resolver, establishing one well-known resolver helped to cope with the rising number of assigning organizations, and the generic N2T design, which works for any kind of identifier, was consistent with the open decentralized ARK philosophy. Since the beginning, N2T stored individual ARKs as well as forwarding rules for the principal persistent identifier schemes (ARK, DOI, Handle, URN, PURL). Over time, N2T has gradually added more forwarding rules, made formal ARK rule replication arrangements with the National Library of France and the US National Library of Medicine, established a database replica in Scotland, and worked with Crossref to load and test resolution of all their DOIs.

However, until recently, N2T.net could not resolve compact identifiers based on the Identifiers.org prefix set, and Identifiers.org could not resolve the prefix-colon-LUI classical compact identifier format. Nor could either system provide a means to declaratively select from among multiple data providers. For example, there are four distinct providers of Worldwide Protein Data Bank (PDB) material, each of with its own presentation format and supplementary material added onto core PDB datasets identified by common accession numbers. Where users need to reference a specific provider’s version, an override of resolver-based default selections is needed.

The JDDCP states, in Principle 4, that “A data citation should include a persistent method for identification that is machine actionable, globally unique, and widely used by a community.” The purpose of this principle is to allow software agents of various kinds to directly resolve a data citation to the underlying landing page, where its metadata resides, and where content negotiation for retrieval of the data itself under access control rules may take place. Data citation without web-resolvable persistent identifiers would be a half-measure blocking development of a fully capable data access ecosystem.

Our view, therefore, is that a harmonized resolution approach for compact identifiers must be available to support biomedical data citation, given the sparsity of DOI implementations among biomedical database providers. To be robust, such an approach requires institutional support at multiple sites and organizations in (at least) both Europe and North America. This requirement motivated our harmonization effort, and the relative importance, stability, robust organizational support, and geographic localization of, Identifiers.org and N2T.net, motivated the selection of these two resolvers for alignment.

## Results

### 1. Harmonized approach for compact identifier resolution

The specific contribution of our work has been to develop and implement a robust, harmonized approach to prefix assignment and resolution in the Identifiers.org and N2T.net meta-resolvers, using a common jointly-managed prefix and provider registry. This is meant to be useful to those seeking to implement direct data citation, by providing common methods in both European-based and North-American-based resolver systems, compatible with the widely-used Compact URI (CURIE)^17,18^ identifier approach; and to serve as a basis for further work in identifier harmonization. We welcome the participation of other parties with stable, long-term institutional support, in providing additional resolvers, and in helping to keep the system globally robust.

The approach we have developed supports reliable resolution in each case, regardless of the original style of LUIs – whether they are bare numeric (e.g. 9606 in NCBI taxonomy), alphanumeric (e.g. 2gc4 in PDB), or whether they have a prefix and colon already when issued by the authority (e.g. MGI:80863 in Mouse Genome Informatics). Examples are shown in Table 1 below.

**Table 1.** Examples of citable meta-resolver-provided persistent URLs and their corresponding default targets.

| Compact ID | Persistent URL | Provider URL (not persistent) | Resolves to |
| --- | --- | --- | --- |
| MGI:80863 | https://identifiers.org/MGI:80863 | https://www.informatics.jax.org/accession/MGI:80863 | JAX Mouse Genome accession number MGI:80863 |
|  | https://n2t.net/MGI:80863 |  |  |
| arrayexpress:E-GEOD-2599 | https://identifiers.org/arrayexpress:E-GEOD-2599 | https://www.ebi.ac.uk/arrayexpress/experiments/E-GEOD-2599 | ArrayExpress accession number E-GEOD-2599 |
|  | https://n2t.net/arrayexpress:E-GEOD-2599 |  |  |
| pdb:2gc4 | https://identifiers.org/pdb:2gc4 | https://www.ebi.ac.uk/pdbe/entry/pdb/2gc4 | Protein Data Bank accession number 2gc4 |
|  | https://n2t.net/pdb:2gc4 |  |  |

### 2. Optional Provider codes

In many cases, a digital entity can be accessed through multiple different locations. For instance, the NCBI Taxonomy^19^, a valuable organism-level classification and nomenclature data resource, is accessible from many sources. As illustrated in Table 2, if we look at a given taxon such as ‘9606’ (human); it can be accessed directly through the primary source (NCBI), or through copies available at the Ontology Lookup Service (OLS)^20–22^ or BioPortal^23,24^. The European Nucleotide Archive also serves NCBI Taxon entities, primarily to organize and access data about the taxon’s corresponding genome assembly. Other examples include the International Nucleotide Sequence Database Collaboration and the Protein Data Bank (PDB)^25^. In many such cases each provider has its own particular virtues in the form of additional services and content.

**Table 2.** Examples of persistent, citable URLs for a single accession (NCBI Taxon 9606), with default and specified providers.

| Compact ID | Provider | Persistent Citable URLs | Resolves to |
| --- | --- | --- | --- |
| taxon:9606 | If not specified, defaults to NCBI | https://identifiers.org/taxon:9606 | NCBI Taxonomy, accession number 9606, NCBI primary |
|  |  | https://n2t.net/taxon:9606 |  |
| taxon:9606 | NCBI (**ncbi**) | [https://identifiers.org/**ncbi**/taxon:9606](https://identifiers.org/<b>ncbi</b>/taxon:9606) | NCBI Taxonomy, accession number 9606, NCBI |
|  |  | [https://n2t.net/**ncbi**/taxon:9606](https://n2t.net/<b>ncbi</b>/taxon:9606) |  |
| taxon:9606 | Ontology Lookup Service (**ols**) | [https://identifiers.org/**ols**/taxon:9606](https://identifiers.org/<b>ols</b>/taxon:9606) | NCBI Taxonomy, accession number 9606, OLS |
|  |  | [https://n2t.net/**ols**/taxon:9606](https://n2t.net/<b>ols</b>/taxon:9606) |  |
| taxon:9606 | NCBO BioPortal (**bptl**) | [https://identifiers.org/**bptl**/taxon:9606](https://identifiers.org/<b>bptl</b>/taxon:9606) | NCBI Taxonomy, accession number 9606, BioPortal |
|  |  | [https://n2t.net/**bptl**/taxon:9606](https://n2t.net/<b>bptl</b>/taxon:9606) |  |

We have therefore added “provider codes” allowing citation at a specific resolver instance and agreed on a set of formal rules for resolution. The provider code is optionally prepended to the repository namespace and separated with a slash (“/”), thus: (<resolver>/)<provider>/<prefix>:<LUI>. Examples are shown in Table 2 below. Where multiple providers exist, and the provider is not specified, the resolver determines where to resolve the request based on its own rules, e.g., taking availability or other criteria into account.

### 3. Rules, Registry and Recommendations

#### Compact Identifiers

A “compact identifier” is a string constructed by concatenating a namespace prefix, a separating colon, and a locally unique identifier (LUI), e.g. pdb:2gc4.

#### Provider Specification

To specify a specific provider, where multiple providers exist, prepend the provider code and a “/” to the compact identifier, e.g. rcsb/pdb:2gc4.

#### Provider Default

Where multiple providers exist, and the provider is not specified in the compact identifier, the resolver will determine where to resolve the request based on its own rules, e.g., taking into account uptime availability, regional preference, or other criteria.

#### Redirect Rule

A URL template associated with the provider code is maintained in the namespace registry, defining how to forward compact identifiers to any specific provider (see 4.2.3 below).

#### Prefix duplication

Some LUIs (e.g., for Gene Ontology records) may contain embedded namespace prefixes. These are ignored where the result of concatenation would be duplication of the prefix and the colon, so a compact identifier example is e.g. GO:0003214, not GO:GO:0003214. However, where prefixes are embedded in accessions without a colon delimiter, e.g. CHEMBL2842, we retain the embedded prefix, thus: chembl.target:CHEMBL2842.

#### Administration

Prefixes and provider codes can be requested by completing a form at https://identifiers.org/request/prefix. Administrators are currently designated EMBL-EBI and CDL staff.

#### Resolution

Resolution of compact identifiers is enabled when they are presented as HTTP URIs by prepending the resolution address, e.g. https://identifiers.org/<compactID> or https://n2t.net/<compactID>.

#### Prefix File

A list of unique namespace prefixes and provider codes in YAML^26,27^ format is available at ark:/13030/c7xk84q2j ^28^. The contents of this file are dedicated to the public domain under the terms of CC0 1.0 Universal ^29^.

### 4. Prefix file

The prefix file consists of a sequence of prefix and provider records. The general elements of a “prefix record” follow.

#### Element: namespace(required)

A string of lowercase letters and digits defining the identifier collection, typically for a given database. The namespace prefix must be unique across the registry.

#### Element: provider(optional)

A string of lowercase letters and digits defining one provider for an identifier namespace prefix. The provider must be unique across the registry.

#### Element: alias(optional)

A string of lowercase letters and digits specifying an alternate name for the namespace. The alias must be unique across the registry.

#### Element: title(required)

A text string containing the full name of the prefix.

#### Element: homepage (optional)

A URL that leads to a web page with more information about the prefix. If the page contains schema.org tags, the meta-resolver may exploit them for descriptive information.

### 5. Implementation

We have implemented this approach in both the Identifiers.org resolver (https://identifiers.org), deployed at the European Molecular Biology Laboratory-European Bioinformatics Institute; and the Name-to-Thing resolver (https://n2t.net), deployed at the California Digital Library. These are longstanding public global identifier resolvers with production-quality implementations. They can be used to support machine-resolvable citation of primary research data, in compliance with funder and science policy recommendations. The new features we describe in this article and their joint maintenance have been agreed in a Memorandum of Understanding between CDL and EMBL-EBI. Their support for compact identifiers formalizes longstanding practice in biomedical informatics and supports recent recommendations on biomedical identifiers^30^ in a long-term sustainable way.

## Discussion

Compact identifiers are a longstanding informal convention in bioinformatics. To be used as globally unique, persistent, web-resolvable identifiers, they require a commonly agreed namespace registry with maintenance rules and clear governance; a set of redirection rules for converting namespace prefixes, provider codes and local identifiers to resolution URLs; and deployed production-quality resolvers with long-term sustainability. An example of formal referencing using a meta-resolver URL with an ArrayExpress reference is shown here:

Aimone, J. B., Leasure, J. L., Perreau, V. M. & Thallmair, M. Transcription profiling of rat spinal cord contusion 35 days after injury (2012). ArrayExpress. https://identifiers.org/arrayexpress:E-GEOD-2599 (2012).

We have extended prior work of the Identifiers.org team at EMBL-EBI in collaboration with the N2T.net and EZID team at the California Digital Library, and other collaborators, to provide these missing elements.

Taken together, we believe that these implemented measures will facilitate data citation in the scholarly literature, as well as data integration, with an initial focus on the Life Sciences domain. New rules flow into an underlying registry hosted and jointly maintained at the EMBL-EBI, and a public version of the registry is available at https://n2t.net/ark:/13030/c7xk84q2j.

Ongoing support for these services is now agreed in a Memorandum of Understanding between the EMBL-EBI and the California Digital Library. We hope they will be of significant assistance to publishers and others concerned with citation and resolution of biomedical data. While not all publishers will be ready immediately to cite data, we believe we have substantially simplified the problem for early adopters.

## Methods

The results published here were developed as part of a larger project, the Data Citation Implementation Pilot (DCIP), a supplement to the NIH-funded bioCADDIE effort^12^. Our work was community-based and organized as a Working Group of FORCE11 (https://force11.org)^31^. DCIP participants were organized into Expert Groups (EGs) focused on biomedical communications ecosystem components (Publishers, Repositories, Identifiers) and project tasks (FAQ Development, Journal Article Tag Suite). The DCIP was led by an international Executive Committee (see Acknowledgements) and was launched at a kickoff meeting held in Boston on February 3, 2016.

This document was the product of the Identifiers Expert Group, which following the kickoff, had bi-weekly conference calls, a face-to-face workshop hosted at Harvard University on June 2, 2016, and numerous interactions via email.

Authors also had in depth face to face discussions at PIDapalooza (Reykjavik, November 9-10, 2016) and at the FORCE11 Summer Institute for Scholarly Communications (La Holla CA, July 31-August 4, 2017).

The approach taken was aimed at harmonizing existing meta-resolvers in a way which (a) was sustainable (b) was consistent with current best-practices (c) provided a bridge to rapid implementation of data citation by biomedical publishers and repositories (d) was as consistent as possible with other efforts such as DataCite and Prefix Commons, without blocking rapid harmonization of the Identifiers.org and N2T.net resolver systems. This approach does have limitations, in that it leaves assignment of local identifiers in the hands of many autonomous repositories. Local repositories therefore retain great influence over how their data is referenced, e.g., how versions are identified. However, our approach has a significant strength in that it allows publishers of biomedical research to meet the JDDCP identifier requirements, without asking all biomedical repositories to re-engineer their systems.

The group made every effort to coordinate with and support the resource indexing work of the bioCADDIE project; a technical representative (Grethe) of that project participated actively and very helpfully in the group, as did Ian Fore, the NIH Science Officer for bioCADDIE. Work was also carefully coordinated with the other DCIP Expert Groups. The senior author of this article participated in the bioCADDIE Executive Committee and provided ongoing progress reports.

Substantial agreement on the approach to be taken was developed in the June 2016 workshop at Harvard, which then enabled software engineering to be undertaken based on these ideas. Continuing work, refinements, and preparation of this article was coordinated based on conference calls.

The group felt it was crucial to build sustainability into the project and therefore a Memorandum of Understanding (MOU) was negotiated between the Directorates of EMBL, EMBL-EBI, and CDL (University of California) as part of this effort, providing for continuing joint support for this work. The MOU provides that EMBL-EBI and CDL will forward the prefixes via their meta-resolvers and that the Namespaces and Provider are jointly determined. The agreement renews automatically, with a prior-notice opt-out option for either party.

A key aspect of sustainability is the organizational persistence of parties to an infrastructure collaboration.

EMBL is Europe’s flagship laboratory for the life sciences, established in 1974 by intergovernmental agreement, registered with the United Nations. EMBL’s organizational basis, as established by this agreement, is independent of, and separate from, participation in the European Union – it is a distinct entity standing on its own. The EMBL-EBI is an outstation of EMBL, located in Hinxton, Cambridgeshire, UK. EMBL-EBI funds and maintains Identifiers.org as core infrastructure for biomedical informatics. Additional funding for development of identifiers.org is provided by ELIXIR, an intergovernmental organization that brings together life science resources from across Europe. Additional funding for specific projects has been provided by the European Commission.

The California Digital Library is a division of the University of California Office of the President. It is the digital library services organization for the University of California system as a whole. CDL manages one of the world’s largest digital research libraries. It is funded and supported by the State of California from tax revenues, as a central resource for its internationally recognized University of California constituent campuses. CDL supports the N2T resolver system as a core service offering.

EMBL-EBI and CDL welcome expressions of interest in joining this collaboration, by other institutions having demonstrable sustainability and a record of providing key persistent services in the global digital science ecosystem. Advice on growing and sustaining the service will be sought from additional experts in the field, and participation in key development projects, such as the European Open Science Cloud and the NIH Data Commons, will be actively promoted.

## Acknowledgements

This work was funded in part by the European Molecular Biology Laboratory (EMBL); the European Commission within the Research Infrastructures program of Horizon 2020, project numbers 654039 (THOR) and 676559 (Excelerate); and by the U.S National Institutes of Health (NIH) as part of the Big Data to Knowledge (BD2K) program, BioCaddie (U24AI117966), and the Monarch Initiative (R24-OD011883), NIH Data Translator OT3 OT-TR-16-001. Work was coordinated by FORCE11 (https://force11.org), a not-for-profit community organization seeking to improve scholarly communication through digital technology.

We would like to thank the Data Citation Implementation Pilot’s Executive Committee members, who helped to guide the overall project direction and provided invaluable strategic advice: Carole Goble, Johanna McEntyre, Joan Starr, Martin Fenner, Simon Hodson, and Chu-nan Hsu.

The authors wish to thank Stephanie Hagstrom of the University of California San Diego and FORCE11 for her extremely helpful administrative and planning work on the Data Citation Implementation Pilot, in organizing workshops and conference calls, and in coordinating website administration. Ian Fore (National Cancer Institute, NIH) participated in our workshops and face-to-face discussions, and provided diligent and thoughtful guidance throughout this effort. Melissa Haendel of Oregon Health and Science University provided several helpful comments on the manuscript.

## Author Contributions

Sarala Wimalaratne and Nick Juty were responsible for all software development and alignment work undertaken on the Identifiers.org system in this project. John Kunze was responsible for development and alignment work undertaken on N2T.net, along with Greg Janée. Their work was critical in achieving the goals of this project, and incorporated many years of experience in engineering and operating their respective resolvers.

Henning Hermjakob provided overall direction for the Identifiers.org component. Rafael Jimenez provided specific architectural guidance from the perspective of ELIXIR EUROPE. Jeffrey Grethe was the key interface of this project with BioCADDIE’s data indexing and search facility development, helped maintain alignment with that work, and provided architectural guidance. Julie McMurry contributed architectural guidance on identifier design incorporating perspectives of the Prefix Commons project.

All authors participated in developing the alignment concepts, evaluating design tradeoffs, matching requirements to implementation, and in writing and/or editing this article.

Wimalaratne, Kunze, McMurry and Clark prepared the draft of this article for the initial submission, with editorial input from their coauthors. Wimalaratne, Kunze, and Clark prepared the final response to reviewers and the revised article.

Maryann Martone and Tim Clark led the Data Citation Implementation Pilot, of which this work forms a component, and ensured that this and all other components, were strategically aligned, properly executed and fully supported each other. Tim Clark organized and led the overall work reported on here, and wrote and edited substantial sections of this article. He had full access to all the data in this study and takes responsibility for the integrity of the work.

